# General Framework for Tracking Neural Activity Over Long-Term Extracellular Recordings

**DOI:** 10.1101/2020.02.11.944686

**Authors:** Fernando Julian Chaure, Hernan Gonzalo Rey

## Abstract

The recent advances in the chronic implantation of electrodes have allowed the collection of extracellular activity from neurons over long periods of time. To fully take advantage of these recordings, it is necessary to track single neurons continuously, particularly when their associated waveform changes over time. Multiple spike sorting algorithms can track drifting neurons but they do not perform well in conditions like a temporary increase in the noise level and changes in the number of detectable neurons. In this work, we present Tracking_Graph, a general framework to track neurons under these conditions. Tracking_Graph can be implemented with different spike sorting algorithms, allowing the experimenter to use the algorithm best fitted for their setup. The main idea behind Tracking_Graph is the blockwise analysis of the recording and application of a classification algorithm to match spikes to templates in different segments, leading to a directional metric that can be used to link clusters across blocks. Moreover, the algorithm can detect and fix sorting errors (splits and merges) in isolated blocks. We compared an implementation of Tracking_Graph with other algorithms using long-term simulations and obtained superior performance in all the metrics.

## 1 Introduction

The recording of extracellular activity from electrodes implanted in the brain is one of the most established techniques in contemporary neuroscience. In the past years, the chronic implantation of electrodes had allowed the collection of data over a long periods of time. Analyzing this data generates new challenges. At first glance, the computational scalability with the amount of data is one of them (Carlson & Carin, 2019), but others related to the changes on the recording’s properties over time are not fully characterized.

One of the main characteristics of a long recording is its stability; with a stable recording we can monitor the same neurons over a long period of time. In general, the stability of the recordings will depend on the electrodes used and the way they are anchored, leading to a large variety of scenarios, with examples such as tetrodes on the dorsal striatum of rats (Schmitzer-Torbert & Redish, 2004), high-density CMOS-integrated microelectrode array on mouse retina (Fiscella et al., 2012), immobile silicon probes in the mouse cortex (Okun et al., 2016), hippocampal multilayer electrode array in moving rats (Senkov et al., 2015), 672 microwires (in arrays with up to 128 wires) on the cortex of macaque monkeys (Nicolelis et al., 2003), independently movable arrays of nichrome electrodes on the macaque dorsolateral prefrontal cortex (Greenberg & Wilson, 2004), up to 1,024 polymer electrodes in freely behaving rats (Chung et al., 2019), Utah arrays into the human neocortex (Mégevand et al., 2017), depth electrodes in the human hippocampus that are anchored to the skull (Rey et al., 2015), and Neuropixels, high-density probes for stable long-term recordings, from single-shank configurations (Jun et al., 2017) to multi-shank versions (Steinmetz et al., 2021), and their successful application to both non-human primates (Trautmann et al., 2023) and humans (Paulk et al., 2022; Chung et al., 2022). These and other recently developed neural recording electrode technologies (Hong & Lieber, 2019) continue to expand our recording capabilities. Furthermore, other experimental factors could affect stability, for example, if the animal has its head fixed, if it is anesthetized, or freely moving.

To fully take advantage of the long-term recordings and study, for example, the variance of neuronal representations (Clopath et al., 2017) or the plasticity in neuronal processing (Lütcke et al., 2013), it is necessary to track neurons even when the stability fluctuates. A clear example of these issues can be found in recording sessions from the human medial temporal lobe where the microelectrodes are inserted inside a flexible probe that is anchored to the skull nearly 60 mm away from the recording site (Rey et al., 2015). In cases like this, similar responses to a given stimulus during sessions run on consecutive days that are associated to putative neurons with a different waveform (as shown in Figure 1) could come from the same neuron following electrode drift, or from different neurons from the same assembly encoding the given stimulus. Tracking the single neuron activity throughout continuous recordings is the only way to discriminate these possibilities.

**Figure 1:**
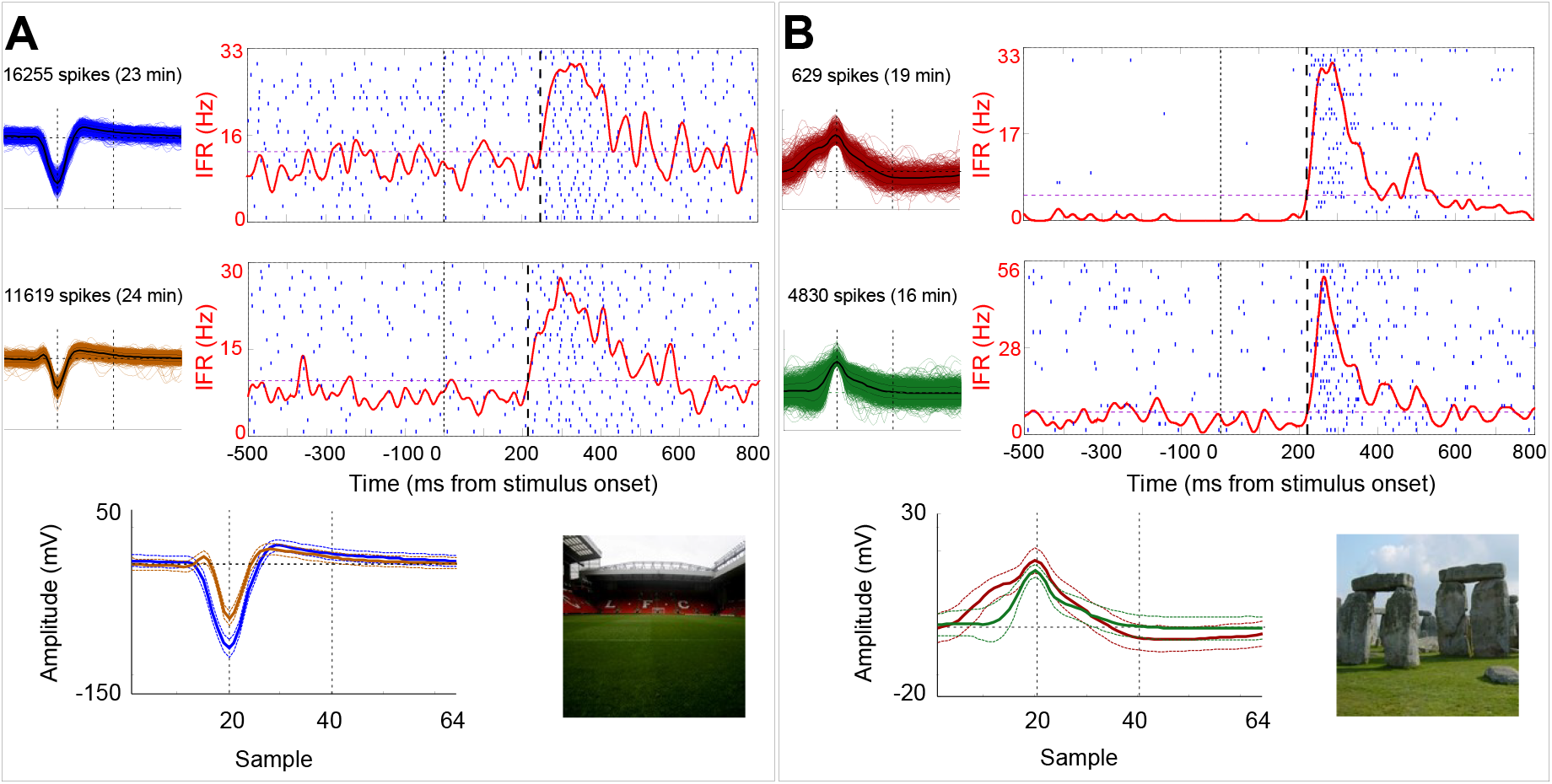
Raster plots of hippocampal neurons recorded from the same microelectrode on consecutive days. **A**: The top panel shows the spikes of a cluster responding to ‘Anfield’ (Liverpool FC stadium) along with its associated raster (each row is a picture presentation, the occurrence of a spike is denoted by a blue line, end the instantaneous firing rate is shown in red). The middle panel presents the results in the following day. Bottom panel shows the spike waveforms compared (mean *±* standard deviation), alongside the image shown to the patient. **B**: Another example from a different patient but with responses to ‘Stonehenge’.

Given the benefits that a tracking method could offer, multiple approaches have been implemented in recent years:

- Performing Spike Sorting on consecutive segments and then combining clusters using a similarity metric (Tolias et al., 2007; Emondi et al., 2004; Niediek et al., 2016). In some cases, additional information such as cross-correlation between neurons is also used (Fraser & Schwartz, 2012). However, this requires neurons with some type of connectivity that allows them to present a statistically distinguishable cross-correlation.
- Associating clusters from discontinuous sessions through waveform similarity and metrics such as inter-spike interval distributions. In these cases, neurons must exhibit particularly stable firing and recording properties, as non-contiguous sessions are not analyzed (Eleryan et al., 2014; Dickey et al., 2009).
- A family of methods has been developed to track neuron drift detected in an initial segment (Franke et al., 2009; Calabrese & Paninski, 2011; Pouzat et al., 2004), but these algorithms are highly sensitive to errors in that first segment and cannot detect the appearance of new neurons after the first segment. These methods emphasize the need for an approach that allows for correction of clustering errors and has sufficient flexibility to detect when neurons are lost below the noise level or when others emerge as the electrode approaches them.

Each of these methods was designed for particular recording conditions, which complicates their application by other groups due to specific parameters and hypotheses. On the other hand, a set of algorithms has approached the problem in a theoretical Bayesian framework:

- In (Bar-Hillel et al., 2006), waveforms are modeled as originating from a mixture of non-stationary Gaussians. First, the recording is divided into small segments, and modeling is performed by optimizing the likelihood while sharing models with neighboring segments. Then, transition probabilities between clusters of each interval are calculated, assuming that each cluster is generated by a neuron whose distribution is a hidden variable. Finally, a global solution is found by optimizing the posterior probability of the model, setting constraints to discard cases where clusters have null transitions in any interval. This work discusses solutions for the random division and combination of clusters generated by the same neuron and supports the detection of a variable number of clusters. However, this method works in two dimensions where Gaussian distributions can be assumed and fails to automatically handle cases such as abrupt changes, sparse neurons, or clustering errors.
- A similar approach is observed in (Wolf & Burdick, 2009), where a causal method is proposed to estimate the Gaussian clusters present in a recording segment, sequentially applying a maximum a posteriori search and using the results from the previous segment as a prior distribution. Therefore, the previous clustering is used as a seed for the next one, while allowing new neurons to appear or disappear below the noise level. The main limitations of this approach are the use of a single predefined space (non-optimal over time), memory limited to the previous segment, and high sensitivity to abrupt changes in the recording.
- The work described in (Shalchyan & Farina, 2014) uses density estimation without assuming the type of cluster distribution in each segment. In this case, previous results are also used to construct a prior distribution for the next segment. Variation in the number of recorded neurons is not considered, and a fixed workspace is taken as the principal components of the first segment, also assuming perfect initial clustering.
- In (Shan et al., 2017), spikes are modeled in a reduced-dimensional space as a mixture of K (value interactively selected by the user) clusters with Student’s t-distribution, whose mean varies in each 1-minute interval window. In this case, a fixed number of neurons is assumed, and maximum likelihood is sought by modeling cluster displacement as a Gaussian with initially estimated covariance. This method requires clusters with constant dispersion (difficult to guarantee in human recordings) and fails to satisfactorily resolve cases where two clusters collide in the workspace.

Although the mentioned methods present good formal approaches, they are not actually used today as they are impractical when analyzing real recordings, due to their high computational cost or lack of publicly available implementation.

In recent years, and due to the rise in the use of probes with specified geometrical contact layout along their structure, a family of methods emerged to estimate and correct drifting. Due to the geometry of these electrodes, drifting is primarily unidirectional, making it possible to estimate it using the geometric relationships between contacts. Then, interpolation is applied to remove the drifting effect on the signal using information from nearby channels (Pachitariu et al., 2024). However, this approach cannot be applied in human recordings with standard depth electrodes, as flexible microelectrodes are used (with skull fixation several centimeters away), with low overlap in the recording volume and whose contacts do not have a fixed position or orientation between them. These recording conditions reinforce the need for a method capable of tracking waveform modifications in each channel independently and allowing for the appearance and disappearance of clusters. Since noise is also dynamic (depending on factors such as the patient’s proximity to their cell phone), temporary merges of clusters and algorithm errors can be expected. For these reasons, it would be ideal to have a visualization tool to monitor what is happening with the isolated clusters during the hours of recording.

The FAST algorithm (Dhawale et al., 2017) is a clear example of the possibility of implementing a method capable of tracking a variable number of neurons over time. This method is based on the successive application of SPC to detect small sub-clusters and a similarity metric based on the distance between sub-clusters. Then, it applies a linear programming algorithm to optimize an objective function, which depends on the distance between means, cluster quality, and multiple constraints. However, despite being a common reference when discussing this type of algorithm, due to its extremely high computational cost, the quantity and lack of interpretability of its parameters, and the difficulty in installing and using it, FAST never managed to be widely adopted by the neuroscience community.

When recording over long periods with drift, each cluster presents its own trajectory in the feature space. Clusters can ‘collide’, meaning they overlap in space at a given moment in such a way that their spikes are not distinguishable. This behavior is observed, for example, when the signal-to-noise ratio (SNR) of two or more nearby clusters decreases (either due to increased noise or electrode displacement); in these cases, the clusters will be indistinguishable while these recording conditions persist. Spike Sorting algorithms that do not use temporal information are sensitive to another, even more common problem, referred to in this work as ‘delayed collision’. This type of collision occurs when two or more clusters occupy the same region of the feature space at any point in their trajectories. The most typical case involves clusters with low-amplitude waveforms and low SNR (i.e., far from the electrode); as the electrode approaches the associated neurons, their spikes will begin to present more distinctive characteristics. However, all neurons that were at a certain distance from the electrode at some point during the recording will have occupied the region of space of low-amplitude and high-dispersion spikes. Algorithms that do not use temporal information might combine all these clusters into the same point cloud, as their clusters overlap in the feature space.

Another algorithm that meets many of the desired requirements is Combinato (Niediek et al., 2016). This algorithm successively applies SPC in each segment and then performs a greedy combination of clusters (within each segment and between different segments) that minimizes distance until it exceeds a fixed threshold. This creates one of its main weaknesses: when combining clusters using only a fixed threshold (associated with how different the waveforms of putative neurons are, as well as the noise level), a fixed separation quality is forced throughout the recording. Another problem is that it does not temporally limit cluster unions, so if two clusters experience a ‘delayed collision’, they will be united even if constant tracking was obtained prior to the union. This creates many problems when analyzing low-amplitude waveforms or low SNR segments. The second drawback of Combinato is its dependence on the clustering algorithm it uses in each segment, which is not necessarily optimal in all cases.

Some Spike Sorting algorithms can tolerate a limited amount of drifting, as long as there are no collisions between waveforms. For example, Mountainsort (Chung et al., 2017) is one of these algorithms. However, the performance of these methods decreases if the recording contains segments with higher noise levels, neurons with low firing rates, or a variable number of neurons (i.e., neurons that appear or disappear below the noise level).

It should be considered that many laboratories have their own Spike Sorting algorithms (published or not) which they trust and whose parameters are optimized for their specific type of recording. This hinders the proliferation of new methods since, even if they can be installed and executed, the parameters would need to be calibrated according to the recording type. This argument reinforces the idea of somehow generalizing currently used methods for non-stationary conditions.

Due to the limitations of current Spike Sorting algorithms and the application difficulties mentioned, we introduce a method called Tracking_Graph to follow neurons over long periods of time and under various stability conditions. The method generalizes Spike Sorting algorithms using simple criteria, modularly separating the clustering problem into segments and solving the transition between segments using a spike classification method. By posing it in this way, it is possible to reduce other algorithms to particular cases, and additionally, a description of the recording in terms of the clusters present over time is obtained, which allows detecting abrupt changes in the recording.

## 2. Proposed Method (Tracking_Graph)

As a first step, Tracking_Graph requires the recording to be segmented into blocks within which a certain degree of stability can be accepted. This approach is used by multiple methods (Eleryan et al., 2014; Dickey et al., 2009; Shalchyan & Farina, 2014; Emondi et al., 2004; Bar-Hillel et al., 2006) and has several advantages, particularly for recording in humans (Niediek et al., 2016). In summary, the benefits of performing a segmentation of the recording are:

- Usually, the high level of noise and/or artifacts particularly affects small time intervals that can easily be confined within a segment. These cuts allow the segments to be made independent and contain the disturbances without affecting the rest of the recording.
- Spike Sorting algorithms scale superlinearly with the number of spikes (in particular, the unsupervised learning component). By separating the recording into segments, the required computation time is limited, and the parallel processing of the first step of the analysis becomes feasible.
- Under certain recording conditions, such as during sleep or experimental sessions, low-firing-rate neurons can increase their firing rate over a short time interval. By analyzing in segments, it is more feasible to successfully isolate these neurons in some of them, as their proportion of events is higher than what would be obtained by analyzing the entire recording.

Ideally, a time interval is sought where classical methods are effective in handling a bounded drift, but which allows the detection of sparse neurons. This parameter will be called T_segment_, which should be chosen by the user as the minimum time interval where the waveforms can be assumed to be stable. For the human recordings covered in this manuscript, a value between 30 and 60 minutes would be reasonable. This contrasts with methods like (Wolf & Burdick, 2009) that require segments on the order of seconds to ensure complete stability. For interpretability and simplicity, segmentation of the recording by time was preferred, as the metric to be considered is how much the waveforms vary over time and the probability of abrupt changes in the recording. Then, it is necessary to apply a Spike Sorting algorithm in each segment (it is not necessary to use the same one in all of them). Once the clusters have been extracted in each segment, an association between them is calculated.

### 2.1 Associating Clusters from Different Segments

It is common for Spike Sorting algorithms to have a stage that applies a classification of this style to assign spikes to known neuron models within the same temporal segment to complement the clustering results (Chaure et al., 2018; Niediek et al., 2016; Chung et al., 2017), or to classify new spikes after an initial training (Eleryan et al., 2014; Franke et al., 2009). This same approach can be used to assign the spikes of a segment B to cluster models detected in segment A. Suppose that the majority of the spikes belonging to cluster B_k_ (cluster k in segment B) are assigned to cluster A_i_ (cluster i in segment A); we denote this correspondence as: B_k_ *→* A_i_. This association is informative, but incomplete, since it could be the case that:

- B_k_ is associated to the same putative neuron as A_i_.
- B_k_ is a spurious split of the cluster A_i_ generated by overclustering.
- A_i_ is a cluster that combines multiple isolated clusters in B, either due to clustering error or increased noise levels.

To solve this ambiguity, the opposite operation can be performed (assigning spikes from A to models from B), which could result in B_k_ *←* A_i_. If this condition is also verified, the notation B_k_*↔*A_i_ is used, and it could be considered that B_k_ and A_i_ come from the same putative neuron in different segments (case 1). If this condition is not met, it is convenient to study what happens with the assignments of the rest of the clusters in B. If another cluster is observed that satisfies B_h_ *→*A_i_, then cases 2 or 3 would be in play. In general, it should be noted that there is not enough information to decide which of these two options is correct without including information from other clusters and segments.

A diagram of the algorithm stages up to this point (segment separation, independent clustering, and association between clusters from different segments) is presented in Figure 2. In panel A, an example of a recording divided into segments is presented, and each one is analyzed by a Spike Sorting algorithm to independently isolate clusters. The example shows an extract of segments S3 and S4, as well as the waveforms of the isolated clusters in each of them (2 clusters identified in S3 and 4 in S4). Panel B presents a two-dimensional schematic of the use of a classifier to estimate the relationships between the clusters of the two segments (S3 and S4). Some spikes are not assigned to any cluster in the other segment due to their long distance; for example, the elements of the purple cluster in segment S4 would not be assigned to any cluster in segment S3. Panel C simplifies the results of the classifiers into a directed graph called the ‘simplified graph’. This graph summarizes the relationships between clusters from different segments in edges that represent which cluster the majority of the spikes of each one would have been assigned to.

**Figure 2:**
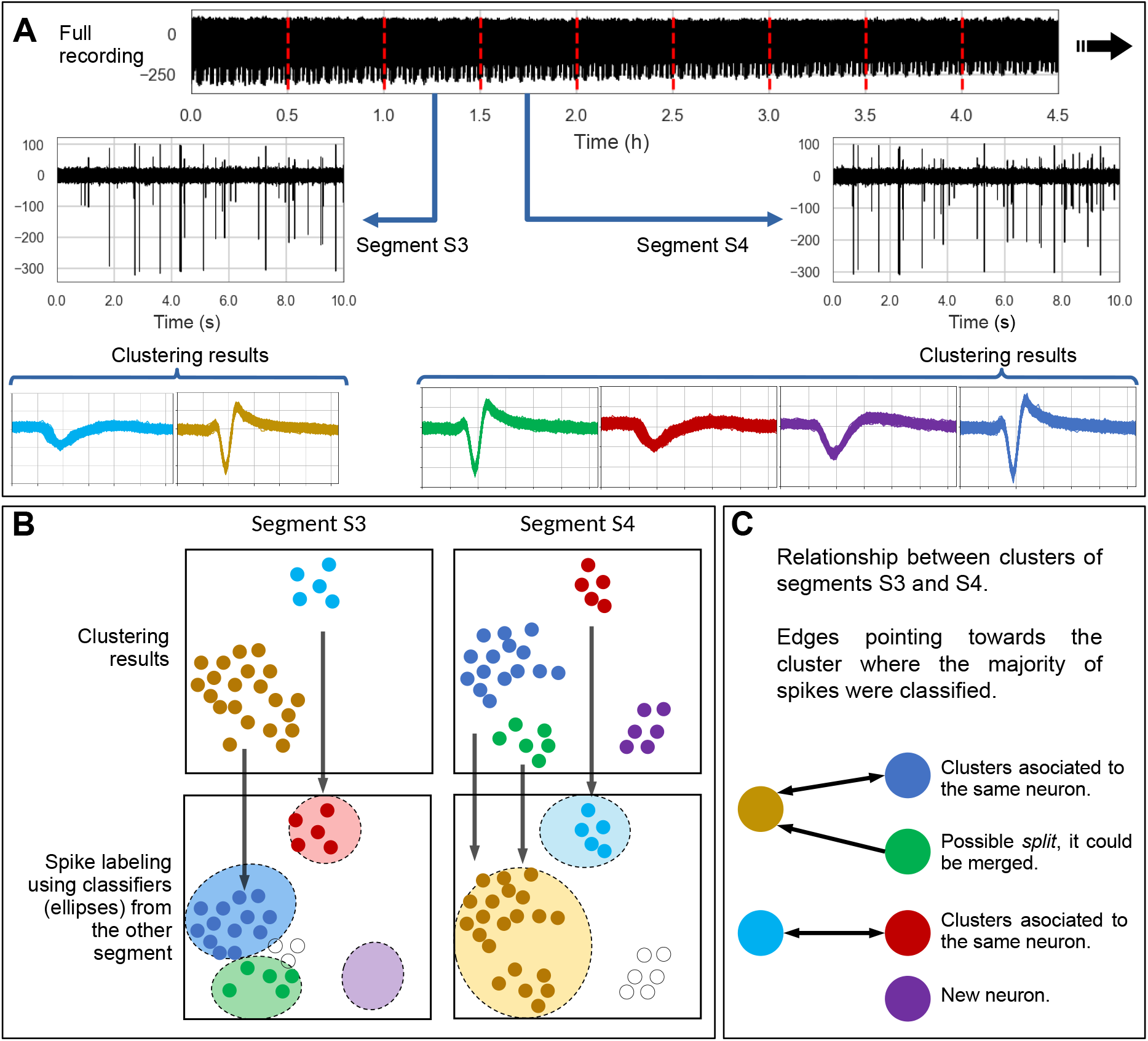
Diagram of the initial stages of Tracking_Graph. **A**: Example of a recording of over 5 hours that is divided into 30-minute segments (red lines mark the segment boundaries). **B**: Example of estimating the relationship between the clusters of two segments, S3 and S4. The upper panels present the spikes in both segments (in two dimensions for simplicity), whose color represents the cluster in which they were grouped when performing independent clustering in each segment. In the second row, these same spikes are presented, but now colored according to how they would be assigned using a classifier (represented by ellipses) trained with the clustering of the other segment. The vertical arrows point to which classifier assigned the majority of the spikes of each original cluster. **C**: Summary of the associations through a graph with a possible explanation of the dynamics of each cluster.

A simple method for performing this classification was used in (Chaure et al., 2018) and consists of assigning each spike to the closest template according to its Euclidean distance, as long as this distance does not exceed a threshold associated with the sum of the variances of the template in each sample. This last detail is important, as it prevents associations with very distant clusters (possibly containing only artifacts). While this classification method was chosen, it is easily generalizable to other distances such as the Mahalanobis distance of each model, or it would even be possible to use a boolean criterion on the distance between the cluster means, assigning a 1 when the distance is less than a threshold. This last criterion is part of the method used by Combinato for tracking. It should be noted that the edges obtained in this way would be bidirectional (due to the symmetry of the distance between means) and, therefore, combine clusters whose means are at a distance less than this threshold. The presented framework highlights the relevant information eliminated by using a criterion of this type, aside from the sensitivity of the results to the value of the used threshold and its complex interpretation in cases of variable SNR.

It is important to highlight the preference for methods that allow partial assignment of spikes from each cluster to multiple clusters from other segments (not just where the majority of spikes go), as they make it possible to quantify how consistent the clustering solutions are with each other, as discussed in Section 1.3. Note that the chosen classification for associating clusters is independent of the Spike Sorting method, presenting a major difference from other works and enabling the combination of diverse methods.

### 2.2 Simplified tracking graph

As observed in Figure 2C, the relationship proposed in the previous section can be represented in a graph, considering each node as a cluster in a specific segment. This graph can be generalized by considering all possible edges between clusters of two segments, where the weight of the directed edges equals the proportion of spikes from the origin that are assigned by the classifier to the destination cluster. A similar approach, but unidirectional and local, is used for monitoring dynamic changes in clusters (Spiliopoulou et al., 2006).

Drifting generates variations in waveforms, and when considering the entire history of neurons in the recording, it would be surprising not to find collisions (particularly if a segment shows high dispersion). Therefore, associating clusters from segments that are very distant in time may lead to false associations. While it is possible to implement more complex criteria, we opted to define a maximum distance in segments L_k_ for relationships between clusters (edges). Both for simulated and real data, we used: L_k_=2, the minimum possible value to allow a cluster not to be isolated in a segment while still maintaining the association chain. In general, the value of L_k_ is associated with the choice of T_segment_, as L_k_ x T_segment_ can be interpreted as the maximum time interval in which a ‘delayed collision’ (quantified in edge weights) would imply that both clusters would have been associated with the same putative neuron. For this reason, a high value of L_k_ would present the drawbacks associated with removing temporal information from collisions, while selecting small values makes it possible for clustering errors (due to decreased SNR or low firing rate) in L_k_-1 segments to prevent the association of clusters that represent the same putative neuron. Here too, we can see a difference from Combinato’s approach, where the criterion for association between clusters has no temporal limit.

Up to this point, the resulting graph, called ‘Generalized Graph’, contains one node per isolated cluster and edges in both directions between all pairs of clusters from different segments (within a temporal neighborhood defined by L_k_). If the only cluster dynamics were merges, splits, appearances, or disappearances, the edges would have weights close to 0 and 1, meaning that all spikes from each cluster would either be classified or not into a cluster from another segment. However, under certain conditions, this may not be the case: for example, a poor choice of assignment method could generate weights far from these extremes, and in practice, cases are observed where automatic clustering presents inconsistencies (for instance, different types of overclustering of the same cluster in consecutive segments), generating relationships that cannot be reduced to simple cases, which implies intermediate values in edge weights. To manually detect and correct these cases, the first step of the graphical interface implemented for the algorithm includes a histogram of algorithm weights and an error matrix to detect segments with complex relationships between clusters (see Section 1.3).

Once cases not described as splits or merges are corrected and accepting some error in edge weights, the graph can be simplified by keeping only edges with weights greater than 0.5 and eliminating the need for weighted edges. This ‘simplified graph’ is sufficient to represent dynamics typically observed in clusters. The example presented in Figure 2C is a simplified graph, so its edges have no weight and do not include information about the proportion of spikes from the brown cluster in segment A that were classified as belonging to the green cluster in segment B. When performing this rounding of edge weights, an error is accepted, which is quantified in the following section.

### 2.3 Detection of inconsistencies

The simplified graph is approximately valid as long as the clustering in each segment can be described in terms of merges, splits, appearances, or disappearances of clusters from adjacent segments. However, due to the high noise level and recording instability, it is necessary to quantify how valid and stable this simplification is for each edge. Since edges are either eliminated or not based on their weight *w*_*a*_(using 0.5 as the cutoff value), it is possible to compute a normalized error for each edge *a* with the following equation:

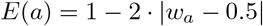

Still, there may be cases where edges have a high error but the simplification does not strongly affect the final result. For example, the combination of two clusters of similar sizes would result in half of the spikes from the combination going to each of its predecessors, generating edges close to 0.5, with error 1. Nevertheless, the edges that would go from the previous clusters to the combined one are close to 1, so they present small errors and would be included in the simplified graph. This way, an edge would always be maintained between the clusters that, depending on the case, would allow them to be combined later. Another equivalent example arises if half of a cluster ‘f’ combines with a larger one ‘g’ in the next segment, generating a high-error edge. However, since ‘g’ is mostly another cluster, this edge would not have its reciprocal, so ‘f’ and ‘g’ would not be automatically associated with the same neuron.

When considering the described examples, it is observed that it is sufficient for one of the two edges between clusters to present a small error to protect the final solution from unstable and/or spurious results. Following this idea, a connection error is defined as the minimum of the errors of the edges connecting two clusters.

Figure 3 shows the connection error matrix for a real data recording in the developed graphical interface, using the notation ‘**S:***{*segment number*}\*|**C:***{*cluster id*}*’. In this example, the connections of one cluster (**S:7**|**C:4**) with two others from the previous segments(**S:6**|**C:4** and **S:6**|**C:6**) show a very high error. This occurred because **S:7**|**C:4** is approximately the combination of half of the other two clusters, a case that cannot be easily reduced to a complete division or combination of clusters. The manual solution for these cases consists of attempting to split **S:7**|**C:4** or merge **S:6**|**C:4** and **S:6**|**C:6**.

**Figure 3:**
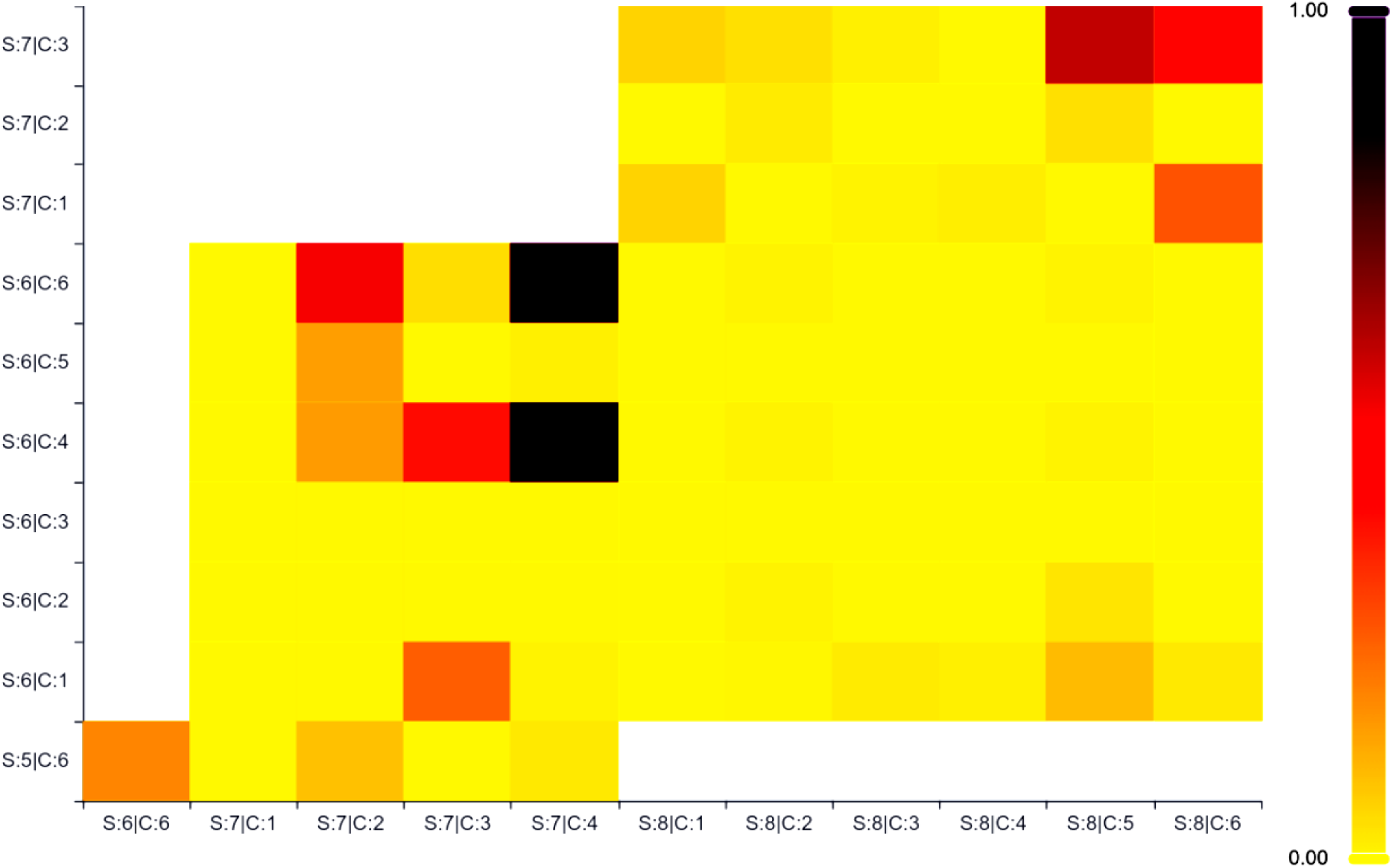
Real data recording where a high connection error was detected that could affect the method’s result. The Tracking_Graph graphical interface reports the connection error between pairs of clusters (using the notation ‘**S:***{*segment number*}\*|**C:***{*cluster id*}*’. The area was enlarged to observe that in column **S:7**|**C:4** (label representing cluster 4 of segment 7) a high value is obtained for rows **S:6**|**C:4** and **S:6**|**C:6** (clusters 4 and 6 of segment 6, respectively).

As this figure is a result of the proposed method, it is useful for detecting problematic segments (or nodes) in which it would be advisable to verify the quality of the Spike Sorting performed. The implemented interface also includes a similar visualization for the error of each edge independently.

### 2.4 Cluster Tracking

From the simplified graph and applying the previously described criterion, pairs of clusters from different segments can be labeled as belonging to the same putative neuron (unit) when they are connected by edges in both directions. It’s worth noting that, given the selected parameters, edges can skip up to one segment, associating clusters that are not temporally adjacent. While this step is useful for tracking clusters over time, in practice many nodes remain isolated or connected to only a few nodes from other segments. Since the objective of this algorithm is to track clusters that can be isolated across multiple segments, the following ‘recombination and pruning’ criterion was applied:

- Stable neurons are detected as those that have nodes in at least N_stable_ segments.
- If a neuron is unstable (with nodes in fewer than N_stable_ segments) but all its edges point to the same putative neuron, the unstable cluster is considered to be overclustering and is merged with that putative neuron. This constitutes a first correction of the sorting using the simplified graph.
- The previous step is repeated until no putative neurons are found that meet these conditions. This resolves cases of overclustering of the same cluster in nearby segments where some edges of an unstable putative neuron can go to both the stable putative neuron and to another one from overclustering.
- Any other unstable putative neuron is discarded and its spikes are labeled as unclassified. The objective of this step is both to discard clusters completely isolated from the rest (e.g., artifacts) as well as clustering algorithm errors that cannot be solved without editing the internal composition of the clusters (e.g., a cluster that is a merge from multiple putative neurons across different segments). A first solution to minimize the number of discarded spikes would be to reassign them to the closest cluster among those in the same segment or neighbors. This proposal is discussed in greater depth in Section 4.1.

The objective of these criteria is to prioritize the tracking of stable neurons, discarding isolated clusters that cannot be tracked over time. The choice of N_stable_ defines the minimum size in segments of the clusters to monitor. While it is associated with T_segment_ to describe it in temporal units, it is also related to L_k_, since if edges are estimated at many segments of distance, the probability of a spurious association of clusters increases, causing the ‘recombination and pruning’ criterion to lose unstable clusters on which to operate. In all the analyses in this manuscript, N_stable_=3 was used.

This criterion aims to be a first application of the simplified graph to solve clustering problems in a segment and isolate artifacts, without requiring the updating of edge weights. The graph obtained after applying the ‘recombination and pruning’ criterion will be called a ‘tracking graph’. In Section 4.1, other possible improvements to this initial approach are discussed. An example of tracking putative neurons is presented in Figure 4, in which cases where the Spike Sorting method presented overclustering were automatically resolved. In the simplified graph, presented in Figure 4A, two nodes (orange and green) do not meet the criterion of having bidirectionally connected nodes in at least 3 segments and are therefore considered unstable. Most spikes from the orange node are always assigned to clusters of the brown putative neuron (outgoing edges always towards brown nodes), but it never absorbs the majority from any other node (no incoming edges), so the criterion considers it a split of that putative neuron in segment 3, to which it will be combined. Different is the case of the green node, whose spikes are assigned to different neurons, and is therefore discarded. With these changes, the final graph is observed in Figure 4B. By having edges that can skip a segment, it was feasible to track the brown cluster even when it was not present in the second segment. This diagram provides useful information about the node in segment 2, which absorbs spikes from both neurons and, although it is mostly contained by the purple neuron in segment 1, can be considered a combination of neurons that could be better discriminated later. In any case, the potential contamination in segment 2 is not preventing the tracking of the purple unit across the other segments.

**Figure 4:**
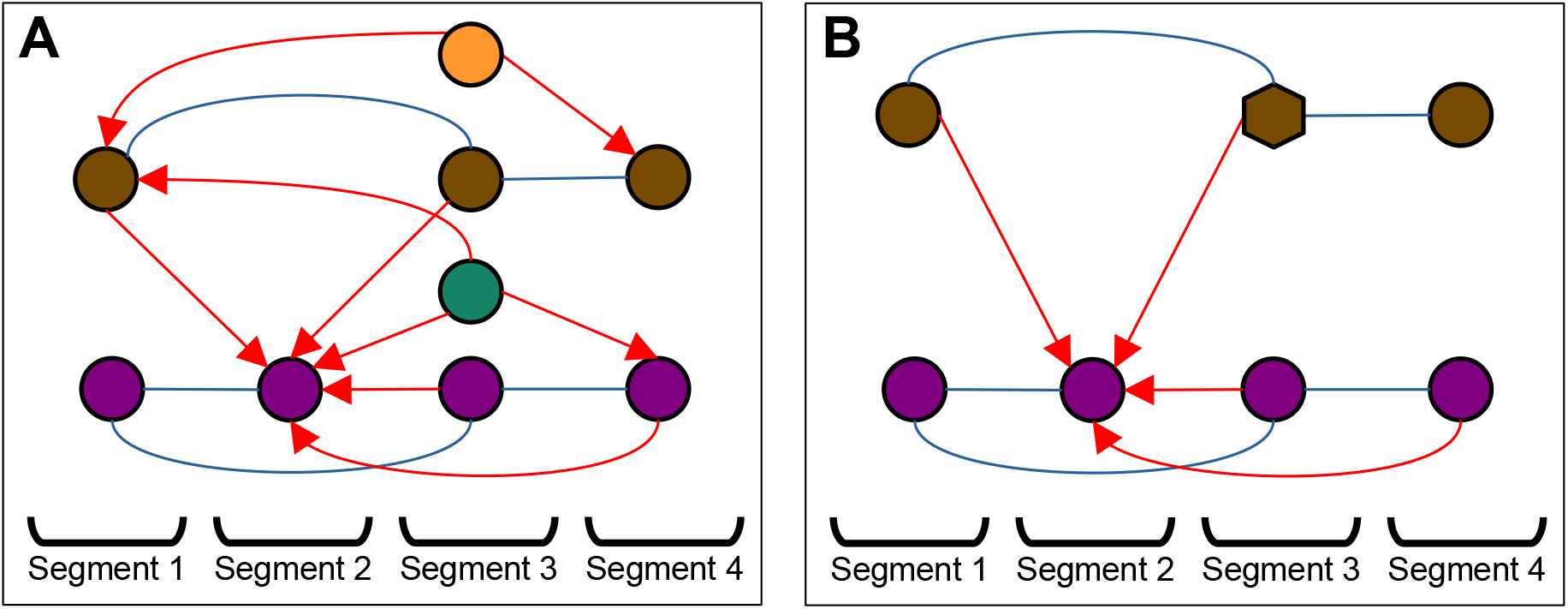
Example of cluster recombination and pruning criterion. **A**: Simplified graph (only edges with weight greater than 0.5 and length less than or equal to two segments) exemplifying the relationship between clusters of a four-segment recording. For simplicity, edges in both directions are plotted as blue lines without arrows, while unidirectional connections between clusters are shown with red lines. Edges between clusters (nodes) of contiguous segments are represented by a straight line, and those that skip a segment are shown with a curved line. line. Nodes were colored to represent association to the same putative neuron prior to the ‘recombination and pruning’ step, as they present bidirectional edges between them. **B**: Resulting graph after simplifying unstable nodes, where the green node was discarded and the orange one was merged with the brown one. A hexagon is used in the brown node of segment 3 to represent that it is not an original single cluster but a combination.

### 2.5 Implementation

The code, implemented in Python, is available in the repository https://github.com/Reylab/tracking_graph. It is compatible with the SpikeInterface toolkbox and designed to allow easy inclusion of new classifiers to define edge weights. It also includes a graphical interface to explore the obtained results.

### 2.6 Visualization and exploration

Selecting the positions of the nodes in the tracking graph to allow for exploration and easy understanding is a challenging task. The first step is to use the position on the x-axis to represent which segment the node belongs to, which does not present any issues; however, the challenge lies in defining the positions on the y-axis, in a way that does not generate long edges that are difficult to follow visually. For example, all the panels in Figure 5 present the same tracking graph with different visual clarity across them. To find a good ordering, classic solutions such as the Fruchterman-Reingold force-directed algorithm cannot be applied, as there is only one dimension to optimize. Additionally, the positions of the nodes on the y-axis are required to meet the following conditions:

**Figure 5:**
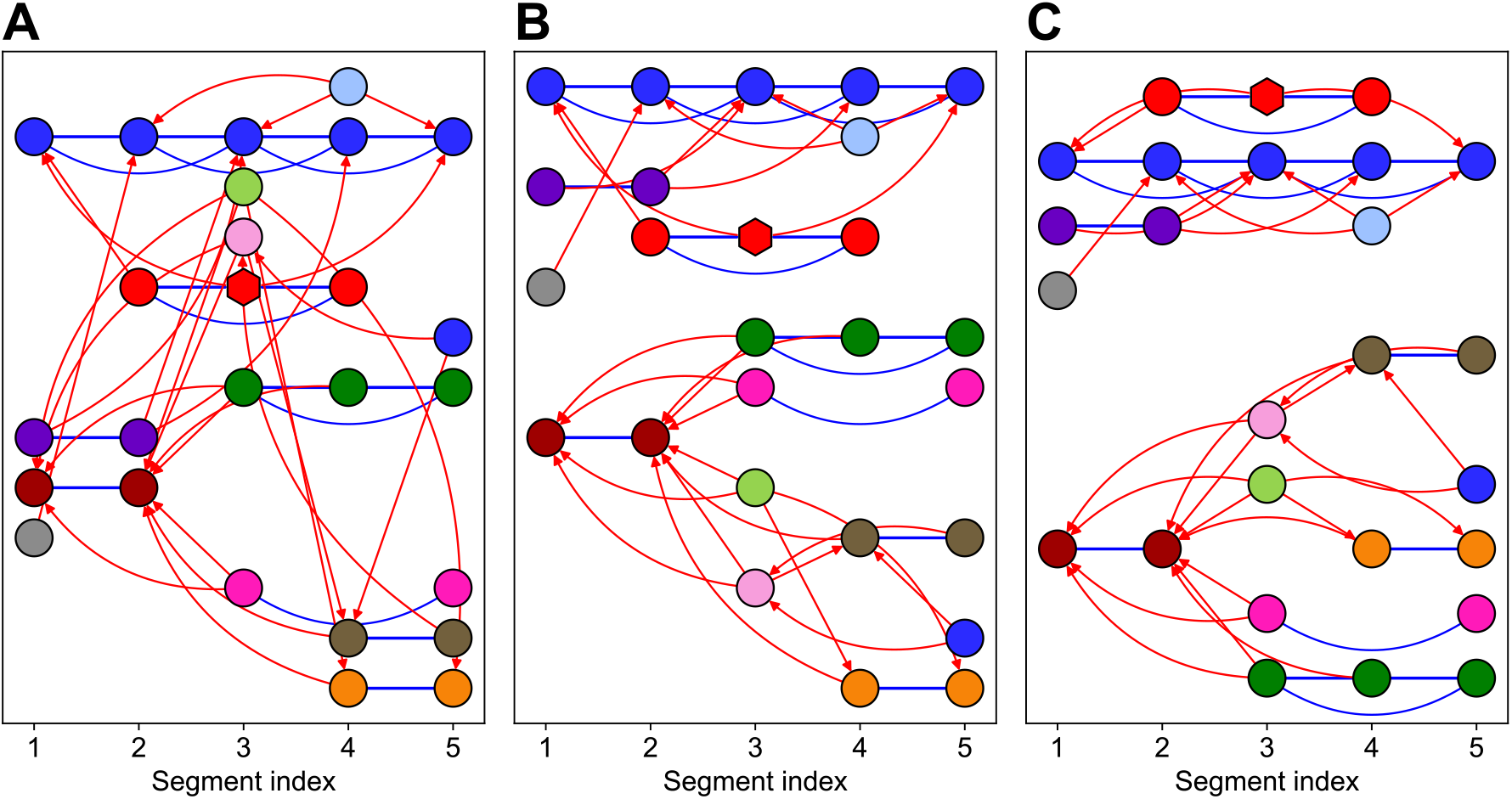
Examples of ordering for a graph with few segments. Bidirectional edges are colored blue and unidirectional ones are red, using straight lines between adjacent segments and curves in other cases. In all cases, the 4 criteria described at the beginning of this section are met. **A**: Random solution with *f*_*loss*_(*Y*) = 755.9. **B**: Minimum cost ordering obtained after generating 5000 possible solutions. This solution achieves an *f*_*loss*_(*Y*) = 178.9. **C**: Solution found by the proposed genetic algorithm after only 10 generations, achieving an *f*_*loss*_(*Y*) = 57.6. Since each generation contains 250 individuals, 2500 values of *f*_*loss*_(*Y*) were required to be computed, half of what was needed for the solution in panel B.

- All nodes associated with the same putative neuron must have the same y-coordinate. This facilitates the detection of segments without the neuron and the time interval in which it was isolated.
- To easily discriminate between nodes and to reinforce their associations, the y-coordinate will be discrete (y *∈* N).
- If a y-coordinate must be reused for more than one putative neuron, there should be no temporal overlap between the putative neurons, and they should be chosen such that there is an empty segment between them, to leave a free x-position and avoid confusion.
- Do not leave empty y-values, to avoid unnecessarily increasing the height of the diagram.

While meeting the previous conditions, the goal is to minimize the length of the edges between groups, primarily reducing the cases where they cross multiple rows. Additionally, as a secondary objective, the aim is to use the smallest possible number of rows, combining rows whenever this does not affect the main objective.

This search for positions can be written as a boolean quadratic programming problem. However, due to the large number of merges and computational complexity, a pragmatic solution was chosen using a genetic algorithm. In this context, the genes are the y-positions of each putative neuron. To simplify and ensure fast convergence, the mutation of a gene/position consists of the following steps:

- A gene to modify is randomly chosen
- Its new position is randomly selected (within the range of used positions and one extra at each end).
- If the resulting position is invalid according to the conditions, the group is randomly placed in a new position above or below the selected one.
- If the initial position becomes empty, all groups are shifted to eliminate the empty space.

Let *dy*_*i,j*_be the distance on the y-axis between groups i and j (note that the same group is always in the same y-position, so *dy*_*i,i*_ = 0); and *n*_*i,j*_ be the number of times that edge is repeated (between the same putative neurons but in different pairs of segments). Thus, the cost function to be minimized is:

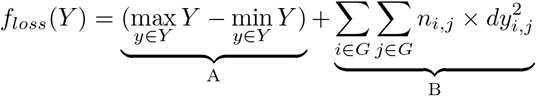

The term A minimizes the total proportion of possible Y values (number of rows) used, but generally contributes less weight than term B, which penalizes the length of the edges in the final graph. The implemented genetic algorithm has the following parameters:

- N_g_: Number of generations (default 300).
- N_p_: Number of individuals in each generation (default 250).
- p_m_: Proportion of genes mutated initially (default 0.3).
- N_s_: Number of individuals that survive from each generation (default 10).

Initially, for each individual, a proportion p_m_ of the positions in its genome are mutated. This amount is reduced linearly so that in the last generation (N_g_) only one position is modified. After each generation, the N_s_ individuals with the lowest *f*_*loss*_(*Y*) are obtained, and these are used to create the next generation. Finally, the chosen positions are those encoded in the individual that minimizes *f*_*loss*_(*Y*) in the last generation.

In panel A of Figure 5 a randomly generated solution is presented, but fulfilling the conditions defined at the beginning of the section. The convention of bidirectional edges in blue color (omitting arrows) and unidirectional in red is maintained, where the edges that connect adjacent clusters of segments are represented with straight lines, while the rest are curvilinear. This example with just a few segments highlights the need to define a cost function *f*_*loss*_(*Y*) and minimize its value to facilitate the interpretability of the results. A brute-force solution to the problem is presented in Figure 5B, for which 5000 random solutions of this type were created, selecting the one with the lowest cost. The solution thus obtained can be more easily interpreted, which is consistent with a value of the cost function 4 times lower. However, proposing solutions of this type do not scale well as the number of possible orderings increases. For this reason, the genetic algorithm presented was implemented, which in 10 generations managed to further reduce the value of the cost function by a factor of 3, computing *f*_*loss*_(*Y*) only half as many times. The solution obtained by the algorithm is shown in Figure 5C, where details can be observed such as: a greater proximity between the blue and red nodes connected by 3 edges, the use of 3 fewer rows, and the fact that the longest edge goes from jumping 3 rows in panel B to jumping only 2 in C.

The algorithm could be implemented using the PyGAD library, which allows the use of a graphics processing unit (GPU) and the addition of techniques typical of this type of algorithms, such as recombination. However, this path would only be necessary in the case of a dynamic version of the graphics and recordings with a large number of channels. The algorithm described in this section is used to order all the graphs presented in Section 3.

## 3 Datasets

### 3.1 Simulations

The dataset was created using a subset of the simulations presented in (Pedreira et al., 2012). From this set, the 15 simulations containing between 4 and 6 neurons were used, a simple scenario for Spike Sorting algorithms under stationary conditions, but with enough neurons so that drifting can generate possible collisions. These simulations were published in (Rey et al., 2015), and have already been used to test Spike Sorting algorithms by other research groups (Niediek et al., 2016). For each synthetic channel, 700 minutes of data were simulated by concatenating original simulations, but instead of using a constant template waveform, we used average spike amplitudes that were linearly increased (or decreased) by a factor of x2.5 (i.e., increased from 1 to 2.5 or decreased from 2.5 to 1), therefore simulating slow electrode drift. In Figure 6A, an exemplary simulation is presented, where the variation of the peak-to-peak amplitude of each neuron can be observed. By varying the waveforms in both directions, more realistic conditions are obtained where each time segment is a different clustering problem and not a rescaled version of the original. It also allows waveforms that were mainly different due to their amplitude to have small collisions, making the separation of the trajectories more difficult. The alignment error was reduced by realigning detected waveforms to the peak, to prevent a disadvantage for algorithms that do not have a stage of this type in the processing pipeline. To prevent too strong similarities between spikes, Gaussian white noise was added, with a standard deviation of 0.25 filtered between 300 Hz and 3 kHz. The noise obtained after this addition was subsequently scaled to maintain the original standard deviation in each recording. This way, the center of the neuron’s cluster was dynamically varied without affecting other high-order statistics when considering small time intervals. A similar approach was used in (Niediek et al., 2016), with the difference that all the spikes were scaled equally. These dynamics are not the expected in a real recording, as when the electrode moves in one direction, some neurons will increase their spike amplitude as they get closer, and others will decrease it as they move away.

**Figure 6:**
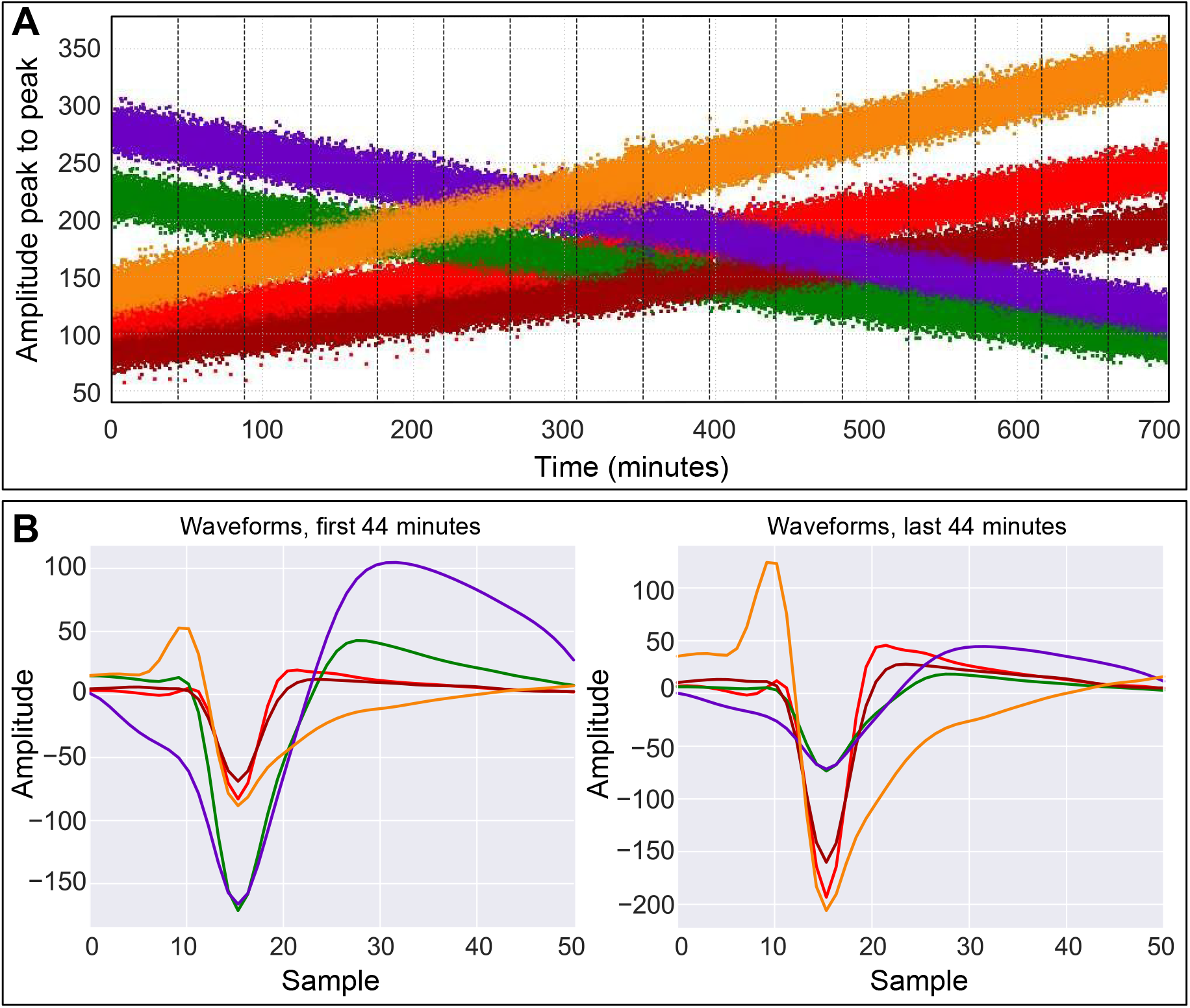
Example of extended simulated recording with 5 neurons. **A**: The peak-to-peak amplitude dynamics of each neuron are shown. At ∼350 minutes, an increase in amplitude dispersion is observed due to the additional noise added. The vertical lines delimit the segments into which the recording was divided for the application of the proposed method. **B**: The mean waveforms of the neurons are shown in the first and last segments of 44 minutes (approximately 20,000 spikes each).

The main drawback of these recordings is the stability of the density of the clusters over time, a characteristic that is uncommon in real recordings where external sources can generate transient increases in the noise level. A typical example can be observed in human recordings when the patient uses a cell phone. As a simpler case, the standard deviation of the noise was doubled in the central 20 minutes of the recording using filtered Gaussian white noise, simulating a transient disturbance.

To perform the analysis using Tracking_Graph, a segment length of 44 minutes was used, to obtain segments similar (on average) to those used by Combinato, which divides the recording into segment of 20,000 spikes. The waveforms of the first and last segments are shown in Figure 6B.

### 3.2 Real recordings

To exemplify the use of the proposed method on real data, we used a recording made at Froedtert Hospital (Milwaukee, USA). In particular, the channel chosen for the example corresponds to a microelectrode located in the left amygdala, and the recording has a total duration of 22 hours.

## 4 Results

### 4.1 Simulations

The simulations were used to verify that Tracking_Graph allows the generalization of different Spike Sorting algorithms under non-stationary conditions.

It is possible to get an idea of the utility of the proposed method by observing Figure 7, where we compared the results of Wave_Clus 3 with Tracking_Graph and Combinato (an algorithm designed to allow tracking of single neurons). As observed in the two upper panels, the proposed method achieves perfect discrimination (even under challenging conditions) and tracked all 6 neurons (plus one additional cluster associated with random threshold crossings of background noise). On the other hand, Combinato (in the lower panels) presented two issues. As observed in the first segment, Combinato’s first issue is its tendency towards overclustering (Chaure et al., 2018). In addition, at times when the waveforms become more similar, Combinato looses tracking, generating new clusters. Its combination criterion does not resolve this issue, and when merging clusters using only Euclidean distance, it generates another issue: the merge of different neurons on certain segments (for example, gray cluster on segments 8, 9 and 14).

**Figure 7:**
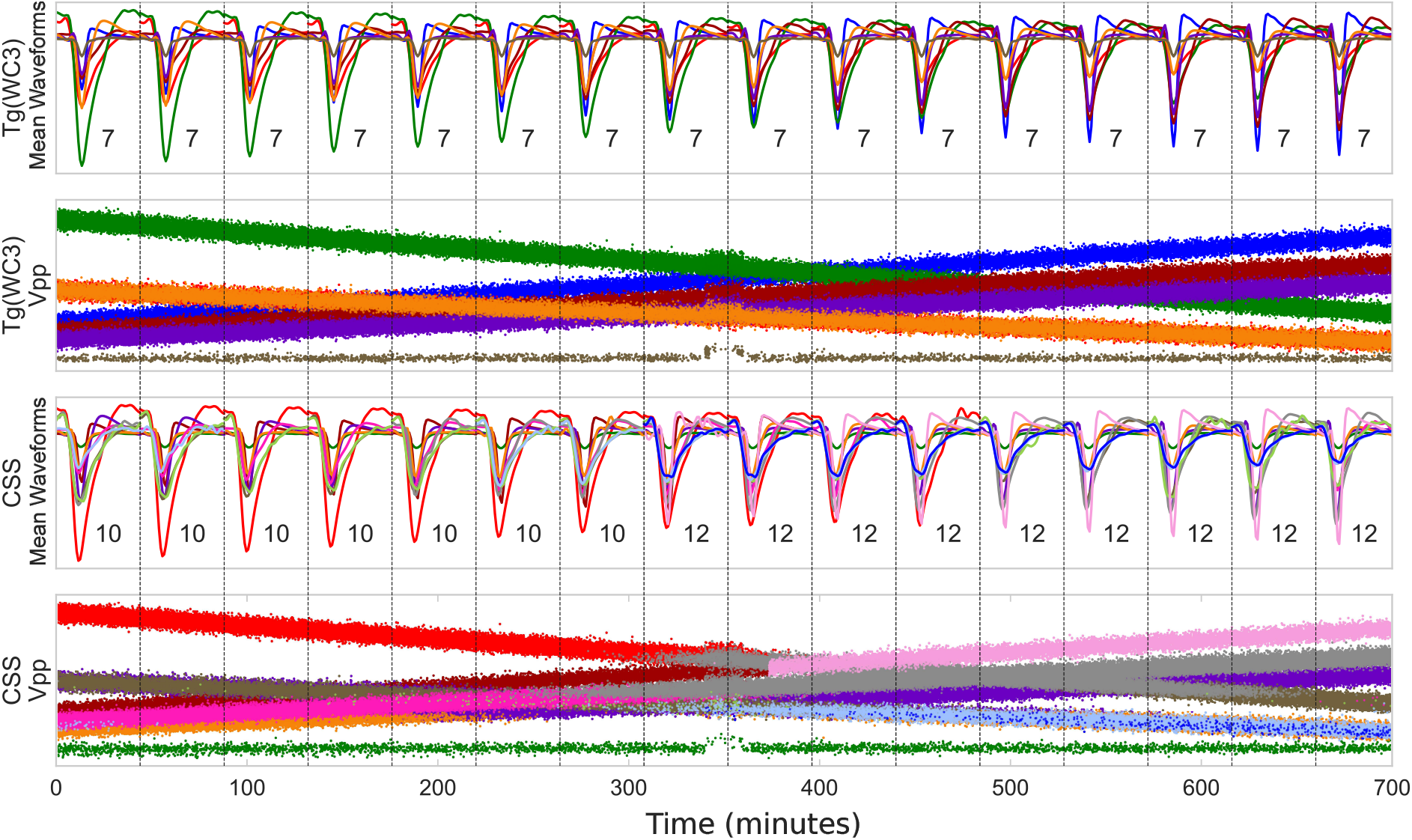
Example of results obtained using Tracking_Graph with Wave_Clus 3 and Combinato alone, an algorithm designed for these types of long-term recordings. For each algorithm, the mean waveforms (and number of detected clusters on each segment) and peak-to-peak value (Vpp) of the spikes are presented. The simulation used contains 6 neurons (3 decreasing their amplitude and the remaining 3 increasing it). The cluster with the lowest amplitude in both solutions consists of spurious background noise detections.

To quantify how Tracking_Graph generalizes Spike Sorting algorithms, the simulations were analyzed using 3 algorithms: Wave_Clus 3 (Chaure et al., 2018), MountainSort (Chung et al., 2017) and Combinato (Niediek et al., 2016). First, the whole recording was processed by each algorithm. Then, it was separated in segments and processed with the addition of Tracking_Graph. In panel A of Figure 8, the Fowlkes–Mallows index is presented for each simulation and method used, quantifying with a single value the similarity between solution and ground truth. This index quantifies how similar two clusterings are (in this case, the ground truth and the solution from each algorithm); if the clusterings are equivalent, the index is 1, and if any pair of clusters across solutions have no pair of element in common, the index is 0. The results show that, when applying Tracking_Graph, solutions closer to the ground truth are obtained. When performing paired sign tests with and without Tracking_Graph, a *p <* 0.001 was obtained for all three algorithms (Wave_Clus 3: *p* = 6.1 *×* 10^*−*5^, MountainSort: *p* = 6.1 *×* 10^*−*5^ and Combinato: *p* = 9.8 *×* 10^*−*4^). In general, Combinato obtains lower indices, which are improved when using Tracking_Graph, but without reaching the values of the other two algorithms. This is because Combinato merges clusters locally, and Tracking_Graph cannot solve this type of errors.

**Figure 8:**
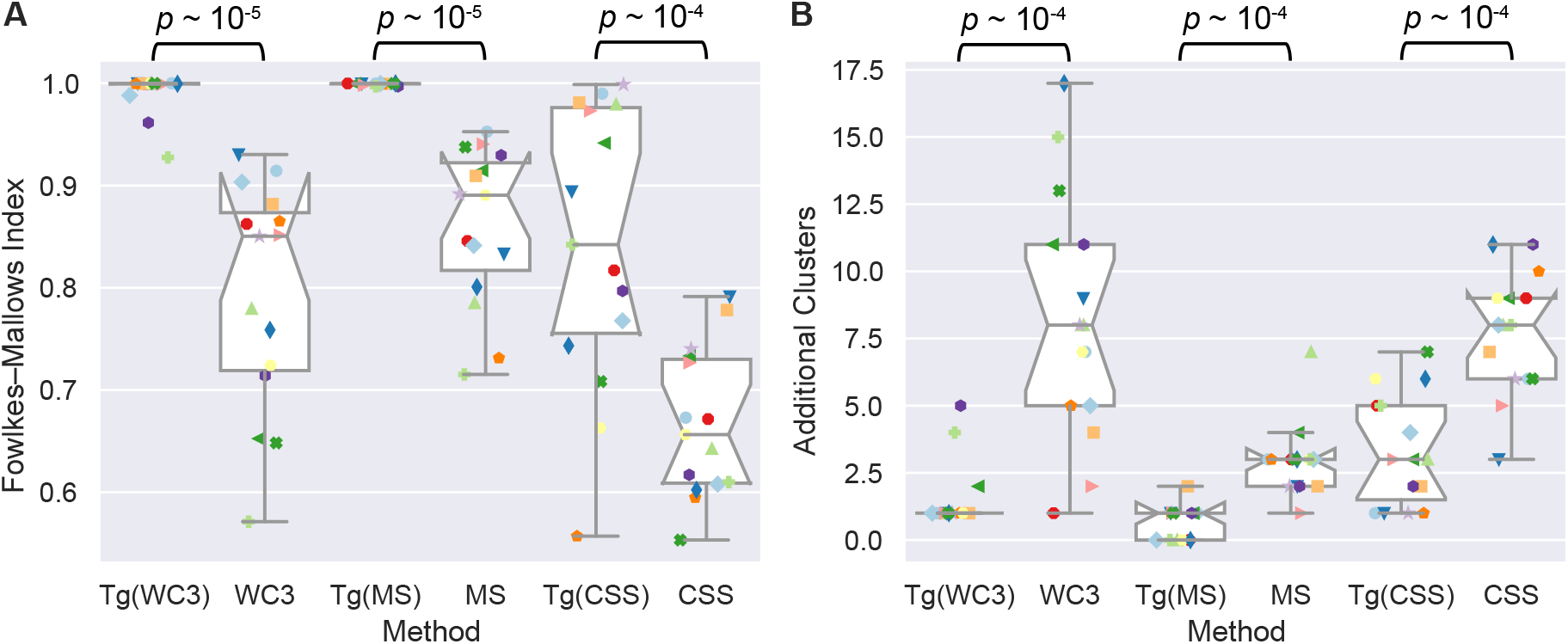
Quantification of sorting performance obtained with simulated recordings. **A**: Fowlkes-Mallows index for solutions obtained using Wave_Clus 3 (WC3), MountainSort (MS), Combinato (CSS) and when applying them with Tracking_Graph (Tg) on the segmented recordings. This index quantifies the degree of similarity between the obtained result and the simulated ground truth. Each symbol is associated to a specific simulated recording. The p-value is associated to a paired sign test between each algorithm and its optimized version using Tracking_Graph. **B**: Number of additional clusters (with respect to the ground truth) for the same recordings analyzed in A.

When processing real data, it is common to find some additional clusters created by overclustering, which are often combined after manual curation. When analyzing long-term recordings, this number could be too high and slow down subsequent analyses, requiring lengthy manual verification. For this reason, we seek to verify if Tracking_Graph manages to decrease the number of additional clusters created. To quantify this, we followed the criterion used in (Buccino et al., 2019), where each simulated neuron obtains an accuracy. This metric is calculated as the maximum of the Jaccard coefficient between the cluster’s activity and each of the simulated neurons. Following this criterion, any cluster with accuracy less than or equal to 0.5 is not associated with any simulated neuron, so it is considered an additional cluster. Through this metric, it is possible to detect if there are cases where the solution is similar to the ground truth but specific neurons are not correctly discriminated. In Figure 8B, the number of additional clusters is presented for each simulation and method. Tracking_Graph significantly reduced the number of addiitional classes for allt he algorithms (Wave_Clus 3: *p* = 1.2 *×* 10^*−*4^, MountainSort: *p* = 2.4 *×* 10^*−*4^ and Combinato: *p* = 9.8 *×* 10^*−*4^).

Using the accuracy metric, it is possible to assign a value to each cluster of the solutions and propose a similar metric, where a value of 1 represents a perfect match between the activity of a cluster and a simulated neuron. In Figure 9, these results are presented, for which an improvement was also observed when applying the proposed approach (Wave_Clus 3: *p* = 5.3 *×* 10^*−*23^, MountainSort: *p* = 2.4 *×* 10^*−*11^ and

**Figure 9:**
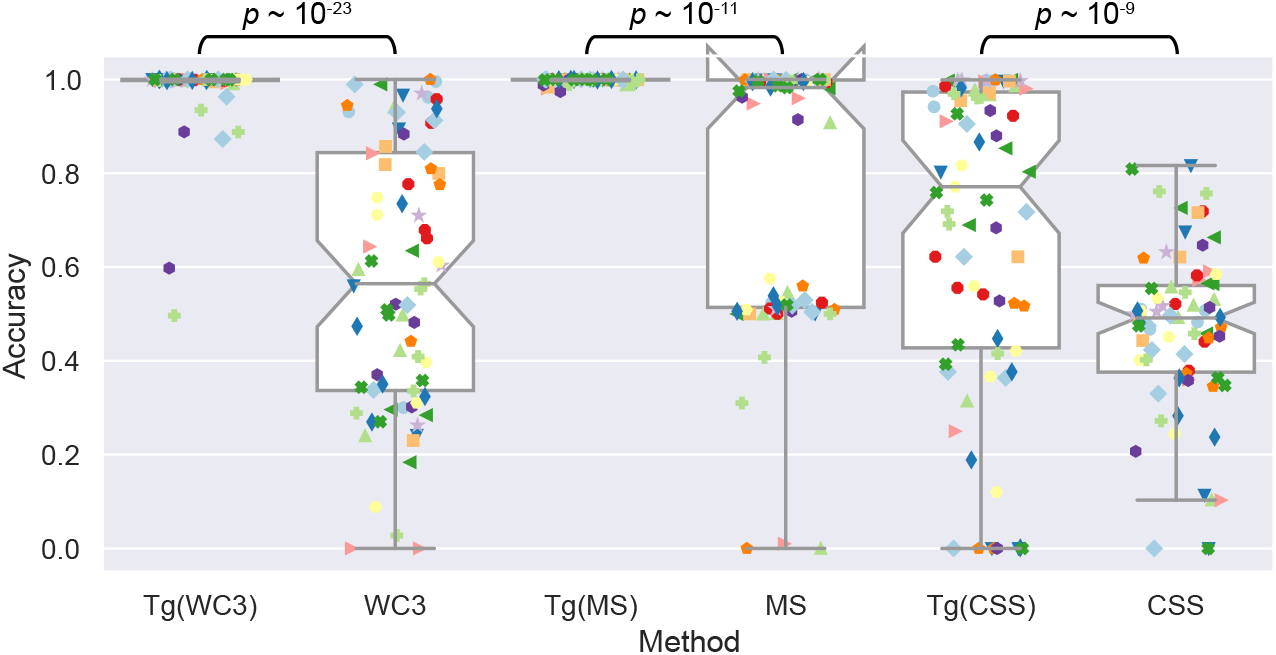
Accuracy of each cluster obtained when analyzing the simulated recordings using different methods. Each symbol represents a simulated cluster on a given recording. As before, Tracking_Graph (Tg) leads to significant performance improvement for all the algorithms.

Combinato: *p* = 1.5 *×* 10^*−*9^). When applying the MountainSort algorithm alone, an accuracy close to 0.5 is obtained in multiple cases, as it fails to overcome the abrupt change in density during the transient increase in noise and ends up dividing the clusters in two.

### 4.2 Exemplary recording from the human amygdala

In this case, each segment consists of one hour of recording, as recordings of this duration have been successfully analyzed previously, and Wave_Clus 3 was used as the Spike Sorting method in each segment. Then, Tracking_Graph was applied to connect the obtained solutions. The same parameters discussed previously were used: edges connecting nodes of up to two consecutive segments and considering clusters as stable if they can be isolated in at least three segments.

In Figure 10, the waveforms of the resulting clusters are presented, as well as the tracking graph. Some clusters can be tracked for multiple hours, although an abrupt change in segment 5 led to new templates that were later tracked for several hours.

**Figure 10:**
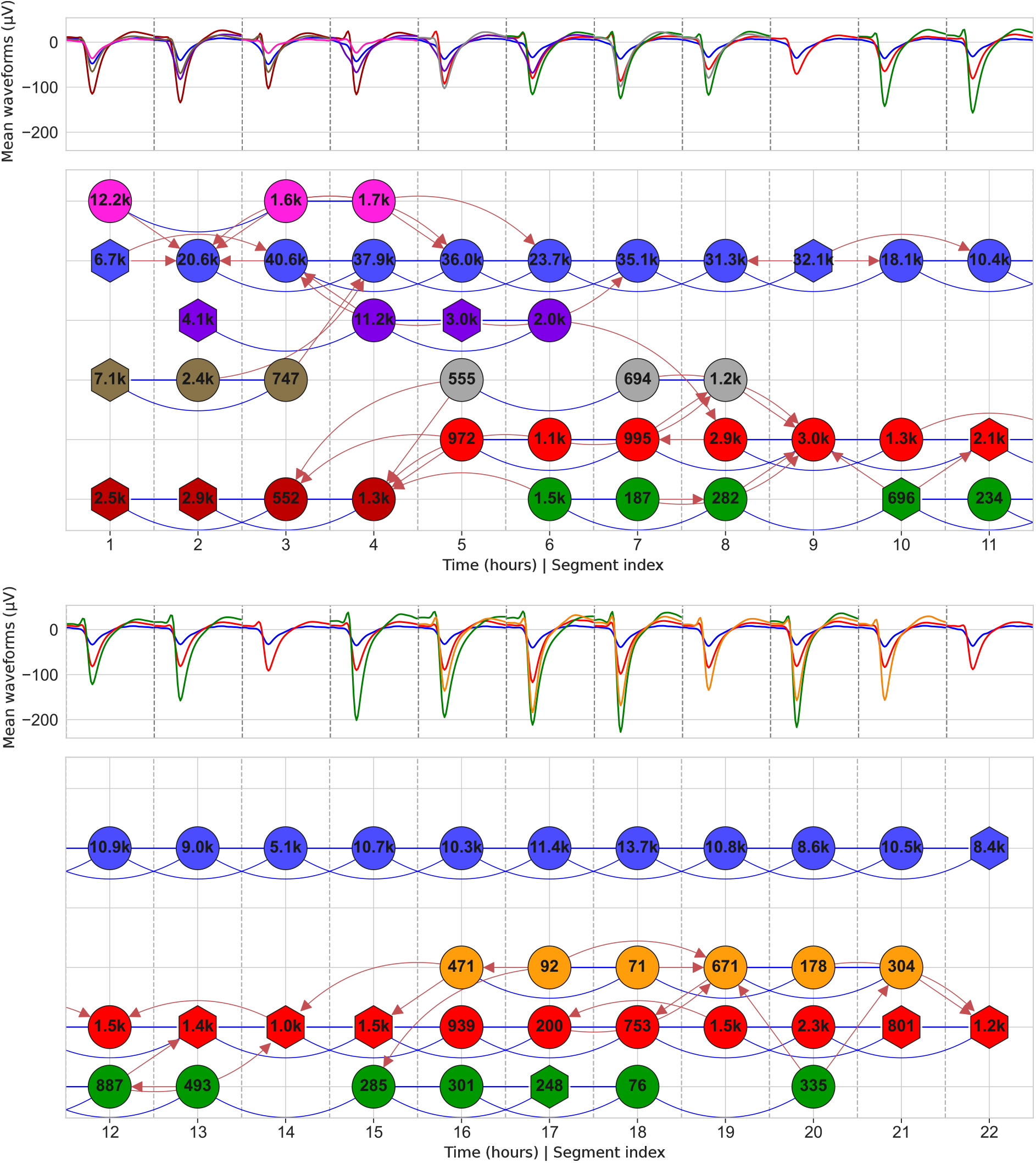
Waveforms and simplified graph of the real recording example. The figure is divided into two main sections, the upper section contains the first 11 hours, and the lower section shows the last 11 hours. The mean waveforms of the nodes in each segment are plotted, where the color indicates which putative neuron they belong to. The lower panel shows the simplified graph obtained by Tracking_Graph. The position of the nodes was calculated using the algorithm described in Section 1.6. The number of spikes is shown inside each node.

In the first segments of Figure 10, it can be observed how the recombination criterion helps containing splits. Wave_Clus 3 can generate overclustering in some solutions by not detecting the end of the superparamagnetic regime, which would explain what is observed in the first segment, where 3 of the 4 nodes are merges of clusters from the local solution. In the first segments, a lower noise level can be inferred, where it is possible to discriminate more stable neurons that eventually dilute into the blue class. This low-amplitude cluster (an unavoidable component of a Spike Sorting solution) is composed of spikes from multiple neurons with amplitude too low to discriminate them, which increases their dispersion, causing the edges of other low-amplitude putative neurons to affect it.

In the second half of the example (Figure 10 lower section), the appearance of a new cluster can be observed in segment 16, with intermediate amplitude between the green and red clusters, which is isolated in multiple segments, making it unlikely that it is due to overclustering of any of these clusters or their partial merge. In segment 14, it can be observed that the green cluster is absent, being combined with the red one, an idea reinforced by the edge that goes from the green node in segment 12 towards that red node. Still, the cluster is restored in segment 15. Similar behavior is observed in the green node of segment 19. These failures in detecting a stable cluster may be due to increased noise, a sorting error, or a low firing rate. In these cases, the simplified graph provides information about the putative neurons present, and can facilitate subsequent correction of the sorting within the segment, as discussed in Section 4.1.

## 5 Discussion

In this work, Tracking_Graph was presented, a tracking and generalization method for Spike Sorting algorithms that require a stationary context. Strong emphasis was placed on the simplicity and interpretability of the results, as well as on the flexibility to use local methods, in order to facilitate its use in other laboratories. In contrast to similar algorithms, Tracking_Graph is easy to apply (without affecting existing analysis pipelines), simple to interpret, and does not require additional hypotheses such as a fixed number of stable neurons.

Real recordings were analyzed, processed by segments with Wave_Clus 3, to observe the characteristics of the results. Through the graph diagram obtained as a secondary result, it is possible to verify the recording conditions that were initially presented such as: variations in waveforms, segments with abrupt changes, waveforms isolated only in some segments, contamination between clusters, etc. Using these graphs and tools, it is possible to estimate the stability and quality of the results, as well as easily detect more stable periods.

As a secondary result, Tracking_Graph allows monitoring and explaining the relationship between clusters in terms of mergers and separations. This descriptive feature brings it closer to a small set of algorithms with this capability, among which it is worth mentioning the general application algorithm described in (Lughofer, 2012), which focuses particularly on solving the problem of joining and separating clusters over time obtained from a data stream.

The Tracking_Graph is being validated using screening results to create a ground truth. In particular, if the same neuron with responses to multiple concepts is detected in more than one session, the goal is to follow that neuron until connecting the clusters of both screenings. Alternatively, neurons that respond to a particular concept could be used, but it would be slightly more likely to be recording a different neuron in each session, which would introduce uncertainties to the ground truth.

### 5.2 Improvements

The presented algorithm prioritizes simplicity and interpretability, leaving several possible improvements to explore:

- **Block Definition:** In this first version, the block was defined as a time interval, but if the goal is to analyze a stable channel with low activity, a definition based on a number of spikes would be more appropriate. Even in general, a hybrid definition could be a compromise solution.
- **Application of other Classification Methods:** In this work, the Wave_Clus spike classification was used, but other more complex methods could be defined. A better metric could have automatically resolved the union of the brown and red clusters in Figure 10. The current code implementation included a classifier based on Mahalanobis distance, but this is only applicable in cases where the number of spikes and the dimensions used allow the calculation of a covariance. This could be solved by working on the principal components of each block or another reduced set of features.
- **Assignment of Missing Spikes:** As a first approximation, the problematic clusters with little temporal presence and without connections or with complex relationships with stable clusters were discarded. A better alternative would be to apply a classification algorithm to assign the spikes to one of the stable clusters with presence in the nearby blocks. However, this step is not trivial, as there are multiple models for each cluster (for example, one block before and in the same block as the spike) and a threshold must be used to continue discarding artifacts.
- **A Posteriori Correction of Spurious Merges:** As observed in several cases, it is normal that even in the simplified graph, there are cases where a cluster presents multiple incoming edges, showing that it is contaminated by more than one stable neuron. In these cases, the classifier could also be used to improve the quality of the clusters, reassigning some spikes to other neurons.
- **Refine the Graph Simplification Step:** If the goal is to limit local errors due to overclustering, the problem of following stable clusters over time could be posed as a Minimum Cost Subgraph Multicut Problem. This type of approach is used to track multiple people in videos when each person can be identified as multiple objects in the same frame (presenting local overclustering) (Tang et al., 2015). In this case, it is important not to fall into the parameter and interpretability problems observed in FAST.
- **Inclusion of Artifact Models:** Under the current design, it would be possible for a classifier to assign ‘spikes’ of artifacts to clusters, contaminating the final results. This can be prevented by including nodes with models that capture non-physiological waveform spikes. However, the characteristics of these models are tied to the type of classifier used. In (Wolf & Burdick, 2009; Bar-Hillel et al., 2006), similar approaches are used to allow spikes to not belong to any neuron model, assigning them to a distribution modeling noise and artifacts.

